# A comparative study of plant phenotyping workflows based on three-dimensional reconstruction from multi-view images

**DOI:** 10.1101/2024.03.21.586185

**Authors:** Daiki Someno, Koji Noshita

**Affiliations:** Department of Biology, Kyushu University, Fukuoka, Fukuoka 819–0395, Japan; Graduate Program of Mathematics for Innovation, Kyushu University, Fukuoka 819– 0395, Japan; Graduate School of Science, Nagoya University, Nagoya 464-8601, Japan; Nara National Research Institute for Cultural Properties, Nara, 630-8002, Japan

**Keywords:** photogrammetry, multi-view image, phenomics, morphometrics, open science

## Abstract

With the world facing escalating food demand, limited agricultural land, and environmental change, there is a growing need for data-driven sustainable agricultural management. Advances in sequencing and sensor networks have reduced costs of acquiring genomic and environmental data; however, collecting phenotypic data, crucial for monitoring plant growth and detecting pests and diseases, remains labor-intensive. Technological advances have enabled efficient collection of three-dimensional (3D) data, yet this process currently involves intricate steps. Therefore, developing effective phenotyping methods is essential. In this study, we developed a phenotyping process based on 3D data, including mask image generation using deep neural network models, 3D reconstruction using the Structure from Motion/Multi-View Stereo (SfM/MVS) pipeline, and surface reconstruction for leaf area estimation. Using soybean datasets, we found that a 1/5.4× magnification effectively generated mask images. Among four mask image usage scenarios in SfM/MVS, applying soybean-and-stage masks before SfM and only soybean masks after SfM yielded the highest-quality point cloud data with the second shortest processing time. Finally, we compared Poisson reconstruction and B-spline surface fitting in leaf area estimation; B-spline fitting showed greater correlation with destructive measurements. We propose an optimal workflow for estimating leaf area and provide tools and datasets for future phenotyping research.

## Introduction

The rise in global food demand ^1,2^, limited agricultural land ^3,4^, and climate change ^5,6^ have heightened the necessity for the quantitative evaluation of crop phenotypic information ^7,8^. This approach would facilitate the development of efficient and robust agricultural management approaches, which are essential for addressing the abovementioned challenges in a data-driven manner ^9,10^. Recent technological breakthroughs, particularly in next-generation sequencing, have facilitated the accumulation of omics data, including genomic and transcriptomic information ^11,12^. In addition, advancements in Internet of Things (IoT) sensor technologies have improved the acquisition of environmental data ^13,14^. However, despite the significance of phenotypic information in understanding plant growth and physiological responses in various environments ^15,16^, the acquisition of phenotypic data remains labor-intensive and yields limited data ^17,18^. Consequently, there has been increasing interest in plant phenotyping technologies among researchers and agricultural producers ^7,8,17–19^. Several studies have reported that the monitoring of three-dimensional morphological properties of plants provides insights for improving cultivation management, including optimizing the timing of pesticide and fertilizer application ^20,21^.

High-resolution 3D morphological data can be acquired efficiently and cost-effectively using 3D laser scanners ^22,23^, depth cameras ^24,25^, and photogrammetry ^26,27^. A pipeline using structure from motion (SfM) ^28,29^ and multi-view stereo (MVS) ^30,31^, which reconstructs a 3D surface as point cloud data from a set of two-dimensional (2D) images captured from different angles, is widely available and implemented in several libraries and software products (e.g., ^28,31,32^). Based on the resultant high-resolution point cloud data, a three-dimensional morphological model can be constructed to yield phenotypic data (e.g., leaf number, plant height, and leaf area index (LAI) ^33,34^).

Optimizing the measurement process for three-dimensional morphological data directly facilitates the efficient acquisition of phenotypic data. Various approaches have been proposed for the acquisition of high-quality three-dimensional point cloud data using the SfM/MVS pipeline (e.g., ^35,36^). Mask images offer several advantages in the SfM/MVS pipeline, such as reduced processing time by limiting the region of interest (ROI) and suppressing noise derived from background textures ^37,38^. Moreover, there are several choices for creating morphological models for estimating phenotypic data from point cloud data acquired using SfM/MVS and 3D laser scanners, depending on the assumptions (e.g., target traits, precision requirements, and data acquisition conditions). Several methods have been proposed to provide robust quantification of leaf morphology, such as modeling the leaf surface as a triangular mesh ^39^, a set of small patches ^40^, and a piecewise polynomial function ^41^. Although Poisson surface reconstruction ^42^ is often used to reconstruct the target surface based on point cloud data in various fields ^43^, it is not ideal for leaf surface reconstruction ^44,45^. This is because it assumes that the target surface is closed (i.e., topologically equivalent to a sphere). Although various techniques for improving the phenotyping processes have been proposed, as noted above, it is difficult to say that methods for optimizing each step in the process or the entire process for the target trait are well established.

In this study, we aimed to optimize the processes for acquiring phenotypic information based on 3D morphological data of plants. Through comparative analyses of various procedures on individual soybean plants, including mask image generation, SfM/MVS pipelines, and leaf surface reconstruction to estimate the leaf area (Fig. 1), we formulated an optimal phenotyping process. To generate mask images using a DNN-based semantic segmentation model, we proposed an ideal scale of input images for the two types of mask images. Within the SfM/MVS pipeline, we compared the processing time and reconstructed point-cloud data quality for the four mask image usage scenarios. Finally, we evaluated the leaf surface reconstruction methods to estimate the area of each leaf by comparing the two general methods. We propose an optimal 3D reconstruction-based plant phenotyping process and provide tools and data to accelerate these comparisons. Such comprehensive comparisons for optimizing plant phenotyping processes can also be applied to other phenotyping procedures. Overall, these optimization processes can yield insights that contribute to the improvement of crop yield, quality, and the cultivation of various target crops.

**Fig 1.**
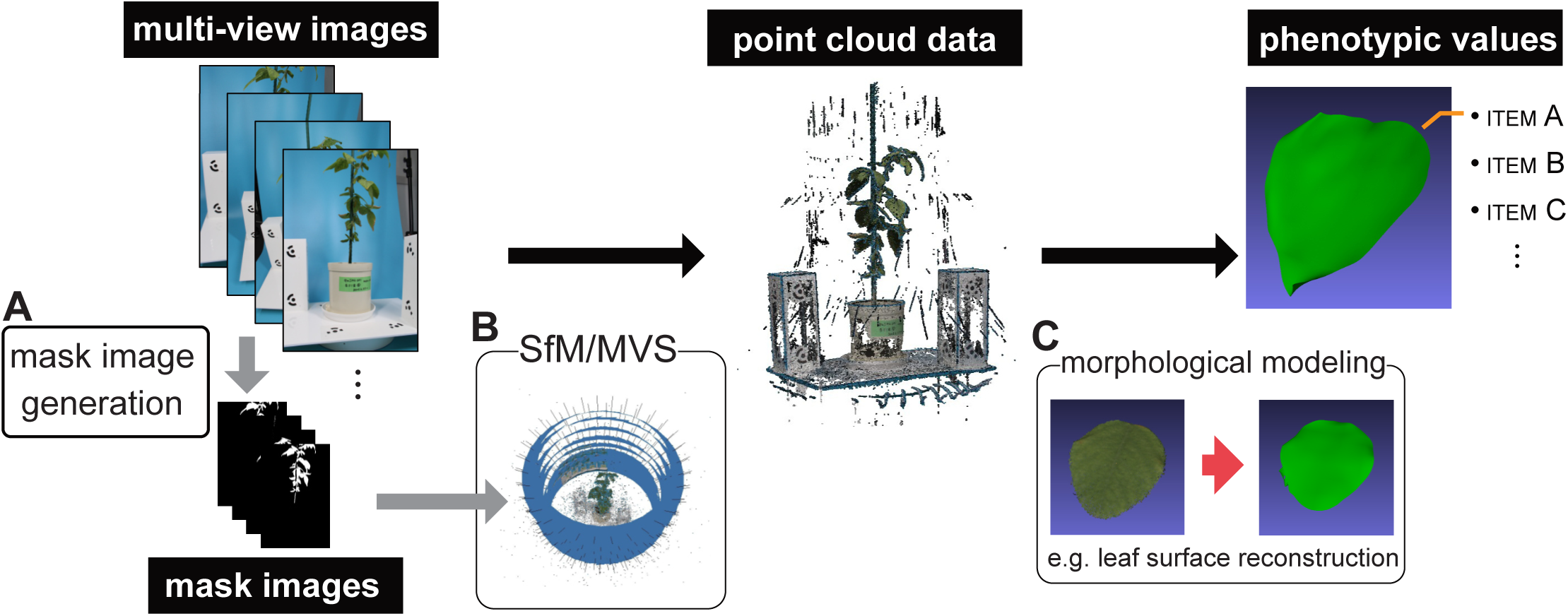
A phenotyping workflow based on the 3D reconstruction. From multi-view images of a target plant individual, mask images are generated for the (A) SfM/MVS pipeline (B). The 3D morphological data is obtained as point cloud data using SfM/MVS pipeline. Several phenotypic values (e.g., leaf area) are estimated through morphological modeling (C).

## Materials and Methods

### Multi-view image datasets of individual soybeans

We used three multi-view image datasets of soybean (*Glycine max*) plants to train the DNN-based semantic segmentation model (dataset_t_), evaluate different SfM/MVS scenarios based on the quality of the reconstructed point cloud data and their processing time (dataset_m_), and evaluate the accuracy of leaf area estimation with two different morphological modeling methods against destructive measurements (dataset_v_). Soybean plants were grown from seeds provided by the National Agriculture and Food Research Organization (NARO) in the greenhouse (25℃) at the Environmental Control Center for Experimental Biology, Kyushu University, Japan.

The dataset_t_ consisted of four individuals from four different cultivars (Enrei, Zairai 51-2, Aoakimame, and Saga zairai), with each individual represented using 256 or 264 multiview images. All images were annotated in a pixel-wise manner across four classes (soybean, stake, stage/pot, and background) using Labelbox (Labelbox, Inc., San Francisco, CA, USA), a web application for generating training data.

The dataset_m_ consisted of six individuals of three cultivars, namely, Seita, N2295, and Keumdu, across different growth stages (4, 8, and 12 weeks after sowing, respectively). The cultivars each included two replicates, with each individual represented using 256, 264, or 272 multi-view images (Table 1).

The dataset_v_ consisted of 256 or 264 images per soybean individual obtained from two cultivars (Hime daizu and Shiro mitsu mame) at two growth stages (6 and 12 weeks after sowing) with two replicates. After acquiring multi-view images, all leaves were sampled from individual plants, and their positions were recorded. The sampled leaves were captured as two-dimensional images at 600 dots per inch (DPI) using a flatbed scanner (CanoScan LiDE 400; Canon, Tokyo, Japan). The region of interest (ROI) for each leaf was set semi-manually using the Selection Brush Tool in Affinity Photo 2 (Serif (Europe) Ltd., Nottingham, UK) to generate mask images for destructive leaf area measurements. These mask images were processed using OpenCV via Python; specifically, this processing involved binarization and the counting of pixels corresponding to each leaf for leaf area measurements. The leaves of Shiro mitsu mame tended to have a smaller area (approximately 1000 mm^2^) than those of Hime daizu (1000–3000 mm^2^).

All multiview images were obtained using a simple photogrammetry studio consisting of a rotating table (MT320RL40, ComXim, Shenzhen, China), eight digital single-lens reflex (DSLR) cameras (EOS Rebel SL1, Canon, Tokyo, Japan), coded markers, and blue screen-like background sheets (Fig. S1). The focal length of each camera was set to 24 mm, and the images were captured approximately every 11° until the rotating table reached 360 °(256, 264, or 272 images per soybean individual).

### Mask image generation using U-Net

U-Net ^46^, a DNN model for semantic segmentation, was used to generate the mask images in the SfM/MVS pipeline. The U-Net was trained for two classification tasks: three-(soybean, stake, and background) and four-class classification (soybean, stake, stage, and background) (Fig 2A). These different classes of mask images can be used across different SfM/MVS scenarios (see **SfM/MVS pipelines** for details).

**Fig 2.**
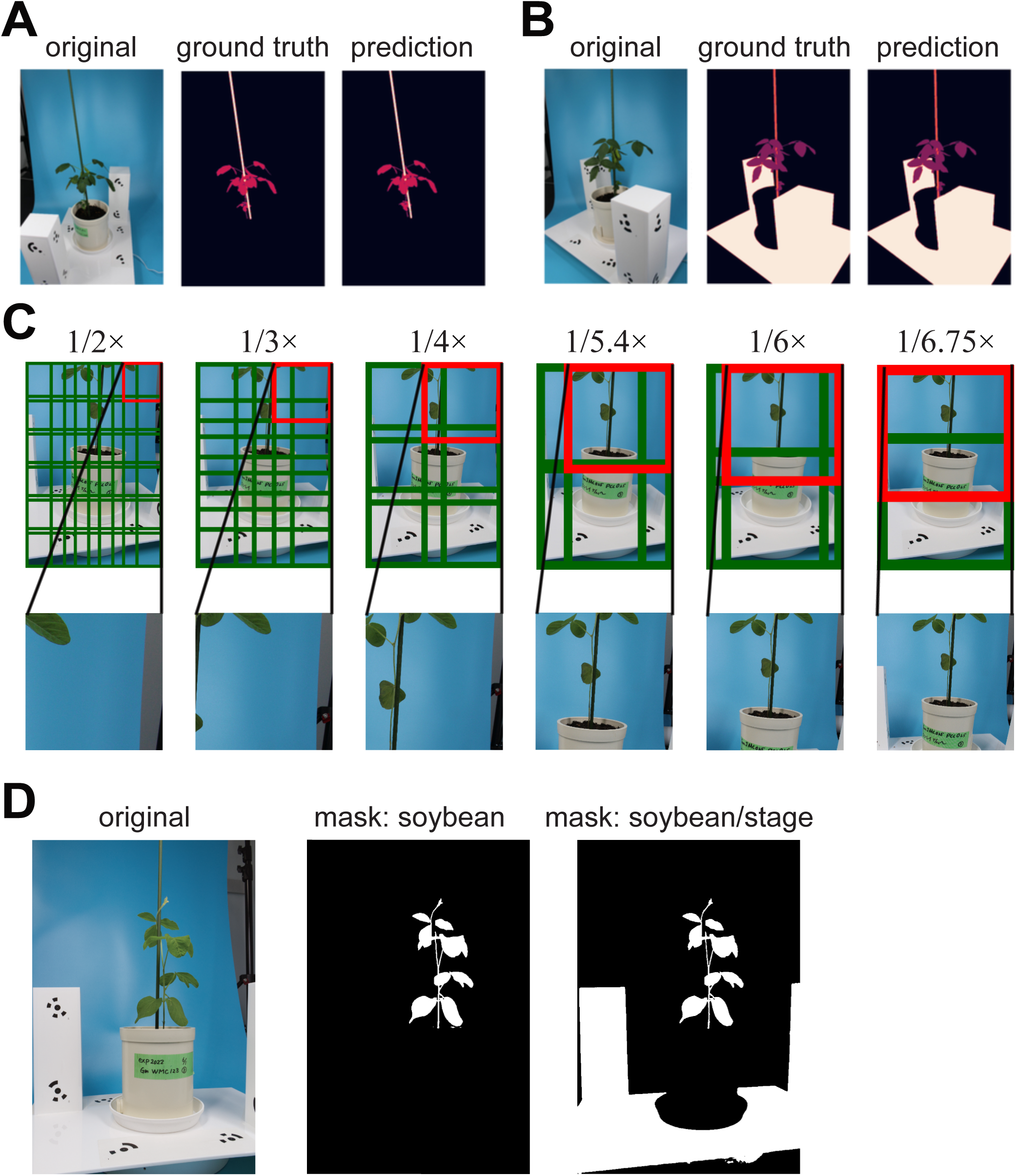
Mask image generation using U-Net model. Results of the semantic segmentation using the U-Net models for (A) three-class segmentation (soybean, stake, and background) and (B) four-class segmentation (soybean, stake, stage, and background) on an individual soybean (Saga zairai). Ground truth data was generated by manual annotation. Each color in the ground truth and the prediction corresponds to each class. (C) Each image in multi-view images was divided and resized into multiple 512×512 tiles with the six different magnifications (1/2×, 1/3×, 1/4×, 1/5.4×, 1/6× and 1/6.75×), for input images to the U-Net models. Lower magnification input image contains larger region (lower row). (D) Soybean mask images (foreground: soybean, background: stake and background) and soybean/stage mask images (foreground: soybean and stage, background: stake and background) were generated based on the predictions of three- and four-class segmentation models, respectively. The examples were mask images of a soybean individual (N2295 #1).

We used U-Net with ResNet34 ^47^ pre-trained on ImageNet as the encoder block and the softmax function as the activation function on the last layer. To train and predict using high-resolution images, each input image was divided into a certain number of tiles, depending on magnification, and resized to 512 × 512 pixels; each tile was segmented into three or four classes and merged into an entire image. An overlap-tile strategy that excluded outer pixels that did not exceed a threshold during merging was adopted to avoid border effects in each tile (Fig. 2B). Class-weighted dice loss was used as a loss function, and we adopted the weights calculated based on the occurrences of the pixels corresponding to each class in the training data (Table 2). The number of epochs is set to 200. Adam, with a learning rate of 0.0001, was initially used as the optimizer; this rate was then changed to 0.00001 after 150 epochs.

Performance, i.e., intersection over union (IoU) and processing time, of U-Net, were evaluated across six different magnifications of the input images (1/2×, 1/3×, 1/4×, 1/5.4×, 1/6× and 1/6.75×) (Fig 3); each image was divided into 30, 15, 6, 4, 4, and 2 tiles on the magnifications 1/2×, 1/3×, 1/4×, 1/5.4×, 1/6× and 1/6.75×, respectively (Fig. 2B). We conducted a group four-fold cross-validation on a dataset_t_; images belonging to a single individual were excluded as test images, and other images were randomly divided into training (80%) and validation (20%) images. In the training step, we used the Python package Albumentations ^48^ for data augmentation, including random resizing and cropping (i.e., while retaining the aspect ratio of the original images), geometrical transformation (i.e., vertical and horizontal flipping, and random rotation by 90° zero or more times), changing color, elastic transformation, and random noise (i.e., Gaussian noising, blurring, motion blurring, and color jittering). The abovementioned steps were conducted using a CPU (Intel(R) Core(TM) i7-6950X CPU @ 3.00GHz) and a GPU GeForce GTX 1080 Ti ×2.

**Fig 3.**
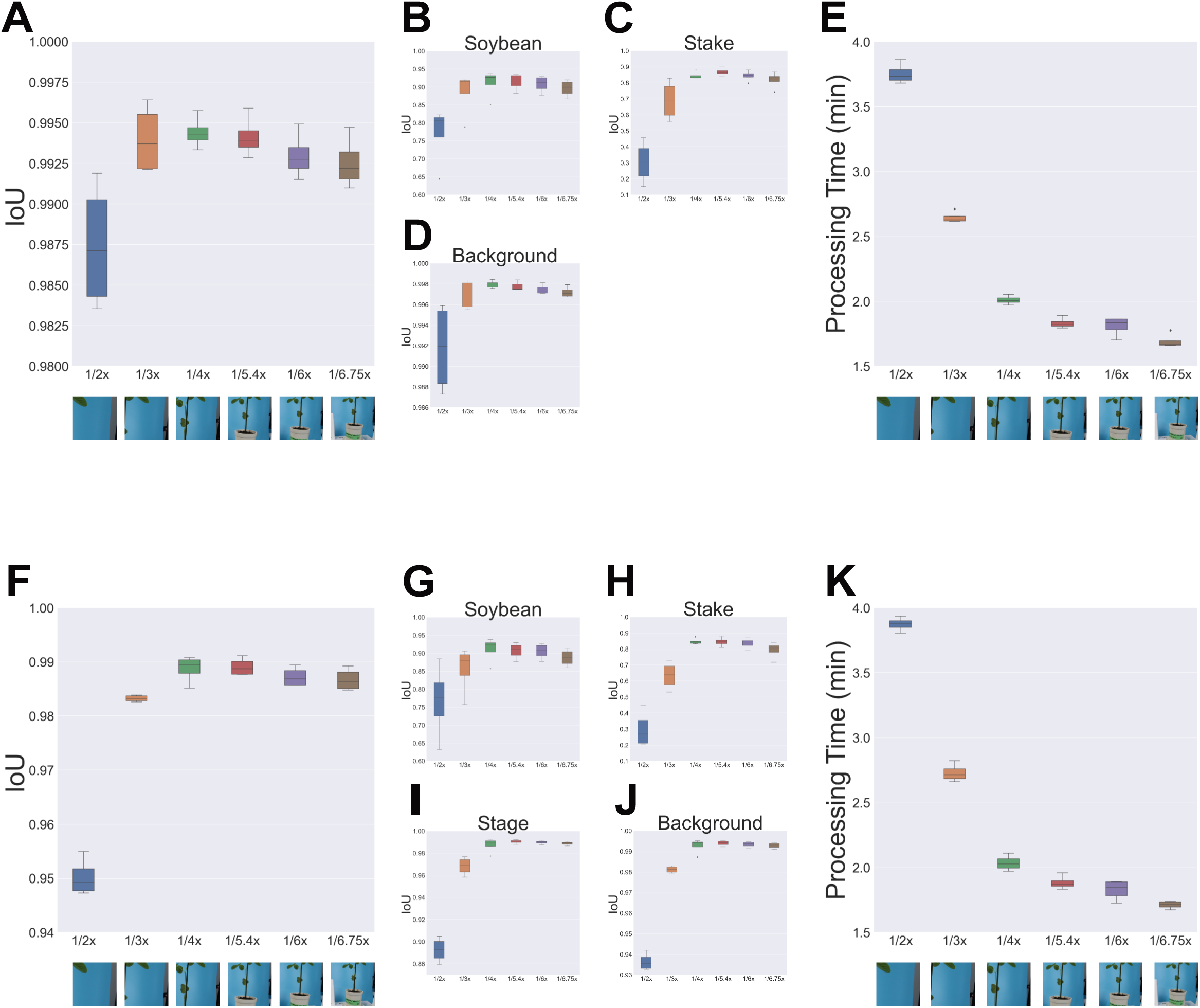
The performance of U-Net models on different magnifications of input image. The IoU scores three-class segmentation for (A) all classes, and each class; (B) soybean, (C) stake, and (D) background. (E) The processing time of three-class segmentation for an individual plant. The IoU scores four-class segmentation for (F) all classes, and each class; (G) soybean, (H) stake, (I) stage, and (J) background. (K) The processing time of four-class segmentation for an individual soybean.

In subsequent analyses, we generated mask images using the U-Net model trained on the entire dataset_t_.

### Web application for generating mask images

We developed a web application for mask generation based on the trained U-Net models (Fig. 4). This application allows users to easily manipulate several parameters (scale, overlap, and mask type) through a simple user interface using Streamlit, which is a Python framework for developing web applications (https://streamlit.io). The web application, operating without a GPU, is hosted on the Streamlit Community Cloud (https://mask-prediction-app.streamlit.app/), and its source code will be made publicly available on the GitHub repository.

**Fig 4.**
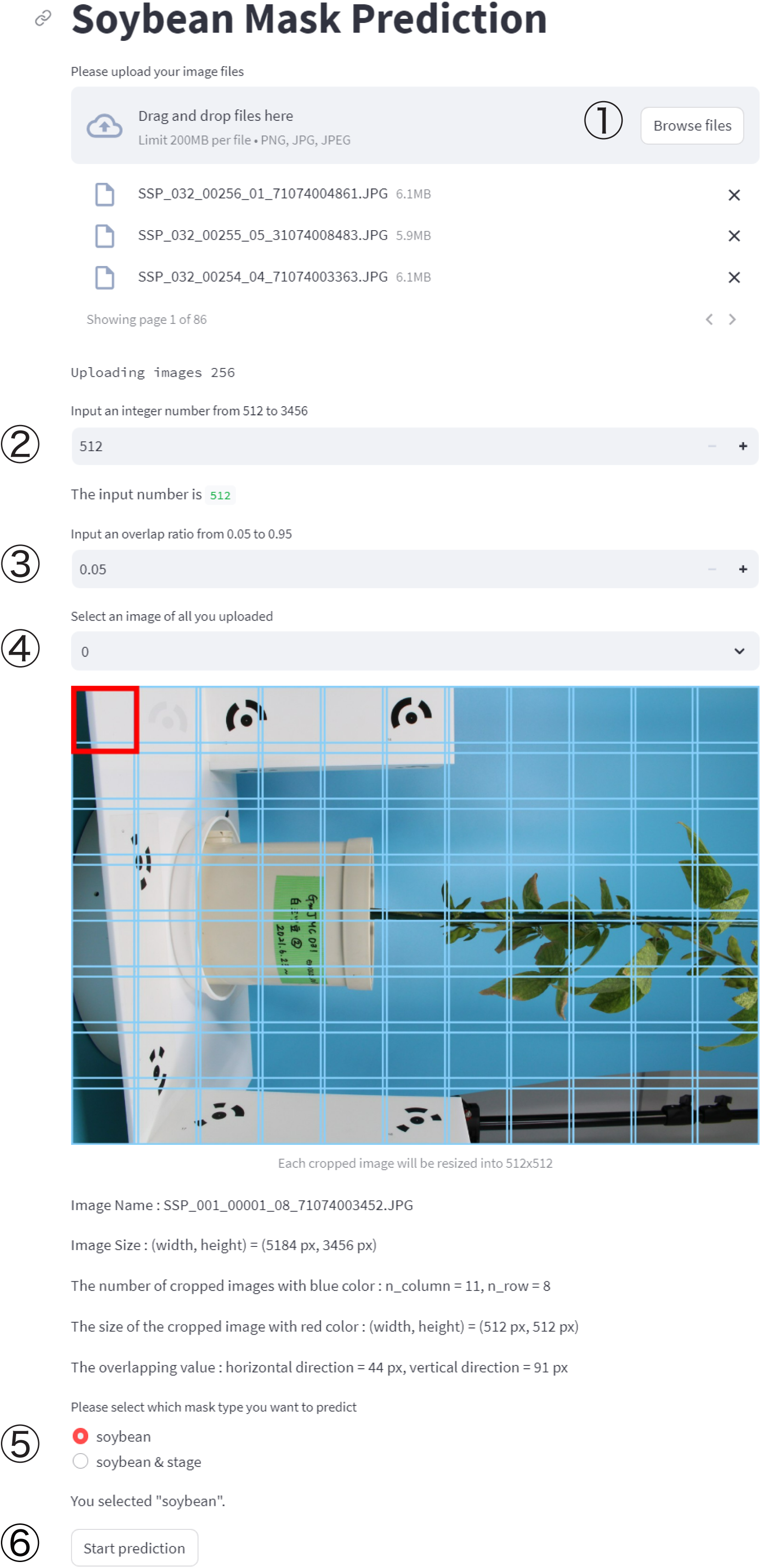
User interface of the web application for mask image generation. (1) Images for generating masks are uploaded via file uploader widget. User sets (2) input image size and (3) overlapping ratio, and (4) selects an image for showing an example for dividing. Finally, (5) one of the mask types is selected, and (6) start prediction. Once processing is complete, the mask images can be downloaded as a zip file.

### SfM/MVS pipelines

We adopted the Structure from Motion (SfM) and Multi-View Stereo (MVS) methods to generate 3D point cloud data. SfM is a photogrammetric method for simultaneously estimating the camera parameters and depth of the corresponding points (i.e., sparse point clouds) from multiview images. The SfM was conducted as per the following steps: (1) key points (characteristic points to be candidates of tie points) were obtained from each image in multi-view images using local features (e.g., SIFT^49^); (2) the tie points, which are corresponding points between images (based on key point matching and outlier removal with epipolar constraints), were selected; (3) an essential matrix for each image pair based on tie points was calculated, and then the extrinsic camera parameters (position, orientation) and the depth of the tie points (sparse point cloud) were estimated; finally, (4) the bundle adjustment was conducted to optimize the extrinsic camera parameters and the depth of the tie points by minimizing the reprojection error. After SfM, the MVS generates dense point-cloud data based on the estimated camera parameters and tie points. Several algorithms have been proposed, e.g., SUrface REconstruction from imagery (SURE)^50^ and patch-based multiview stereo (PMVS) ^51^. We used Metashape v1.8.4 (Agisoft, St. Petersburg, Russia), to conduct the SfM/MVS pipeline. Metashape adopts a SURE-like algorithm to implement the MVS. SURE can offer a depth map, which is the distance from each pixel in a 2D image to the object or scene captured in each image using a semiglobal matching (SGM) -based algorithm^52^, and convert the depth maps into 3D coordinates using a fusion method based on geometric constraints ^53^.

We evaluated four scenarios based on SfM/MVS: (i) no mask, (ii) masks before SfM, (iii) masks after SfM, and (iv) two masks (Fig 5). The basic procedure for the SfM/MVS implementation for each scenario was as follows: (1) multi-view images of a soybean individual were imported, (2) the coded markers were detected, (3) scale bars were created from the pairs of detected coded markers, (4) sparse point clouds were generated based on SfM, with the maximum number of key points set to 200,000 and the maximum number of tie points set to 80,000 in this study, (5) a depth map was created, and (6) dense point clouds were generated based on MVS. In scenarios (ii) and (iii), the soybean mask images were imported before and after step (4), respectively. In scenario (iv), the soybean/stage mask images were imported before step (4) and replaced with the soybean mask images before step (6). These four scenarios were implemented using the Metashape Python API.

**Fig 5.**
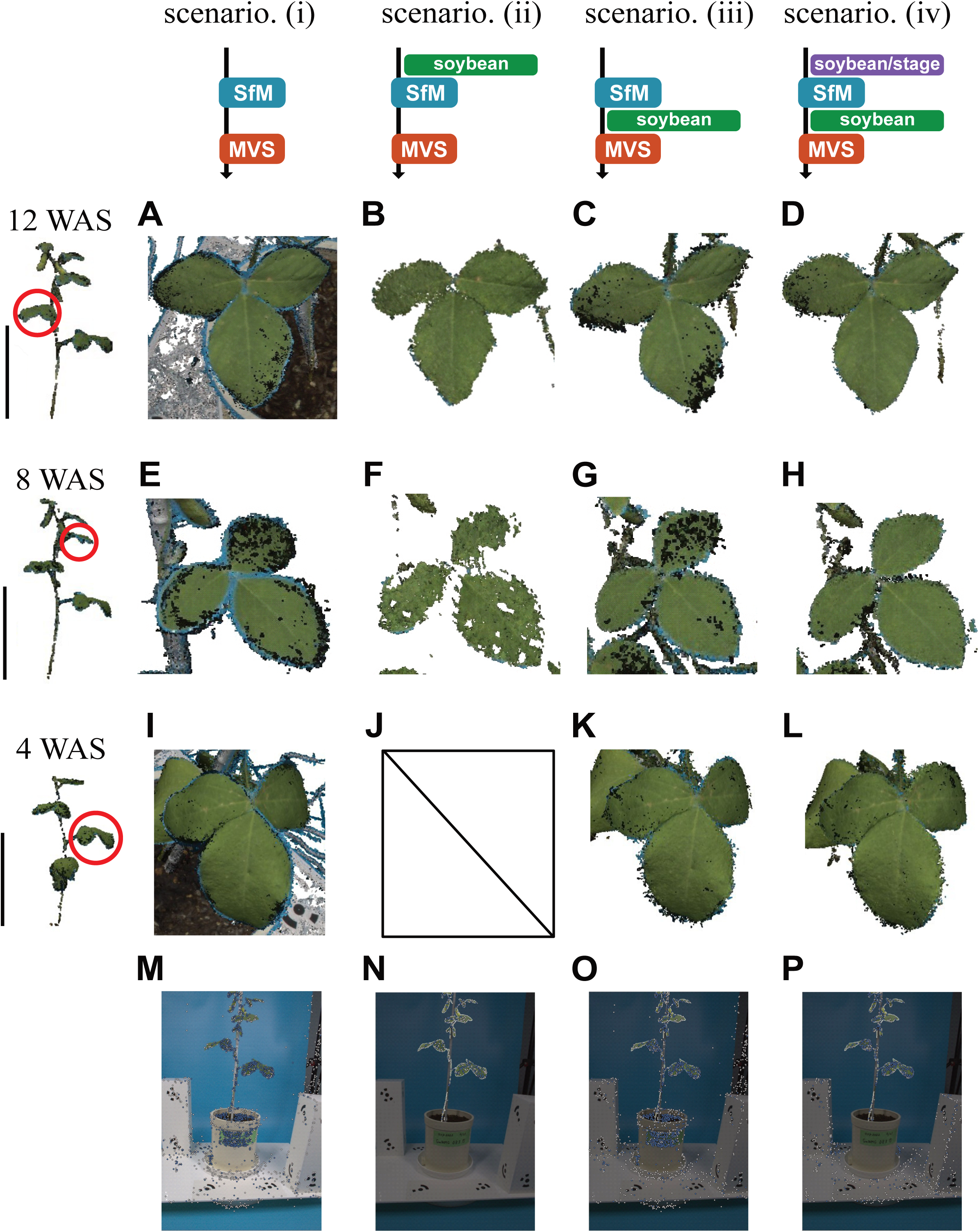
The 3D reconstruction based on SfM/MVS pipeline with the four different mask usage scenarios. Each column corresponds to (i) no masks, (ii) masks before SfM, (iii) masks after SfM, and (iv) two masks. The reconstructed point clouds of a soybean individual (Keumdu #1) at 12 WAS (the second row), 8 WAS (the third row), and 4 WAS (the fourth row). The black bars represent 20 cm. (A-L) Leaflets corresponding to red circles in the first left column. (M-P) Key points and tie points detected in the four different scenarios.

In addition, the abovementioned four scenarios were evaluated for each soybean individual of dataset_m_. In the implemented script, the processing time for each step was measured using the time function in the time module of the standard Python library. The Mann-Whitney *U* test with Bonferroni correction was subsequently performed to investigate the differences in processing times among the four scenarios.

### Leaf surface reconstruction

To estimate phenotypic values from the reconstructed point-cloud data, we employed two prevalent methods for leaf surface reconstruction: (1) Poisson surface reconstruction and (2) B-spline surface fitting.

Poisson surface reconstruction ^42^ is widely used to estimate surfaces from point-cloud data using their normal vectors (i.e., oriented points). Assume that we will reconstruct the boundary 𝜕𝐴 of a target structure *A* from the given oriented points. If a function (called *indicator function* 𝜒), which returns one if the point is located inside of the *A*, otherwise 0, is given, the boundary is approximated by its isosurface. The best-fitted boundary will be estimated by minimizing the differences between the gradient of the indicator function ∇𝜒 and the vector field defined by the oriented points 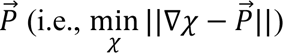. To solve the problem, this equation is transformed into the Poisson equation as follows: 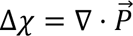. Finally, the isosurface is extracted from the estimated 𝜒. Additionally, the reconstructed surface is filtered based on the point density (i.e., only parts of the surface supported by a sufficient number of points remain) because the method potentially reconstructs a surface in which points are distributed with low density, including the outer region of the leaf surface (i.e., extrapolation). In this study, we used the create_from_point_cloud_poisson function implemented in Open3D^54^ to estimate the leaf surface from the point-cloud data using Poisson surface reconstruction. Vertices of the reconstructed surface were filtered if less than 6.5 points supported them.

B-spline surface fitting is widely used for surface reconstruction from point-cloud data. In this study, we adopted the B-spline surface fitting and B-spline curve trimming approaches ^55^ to reconstruct the leaf surface. A B-spline basis function of degree *d* is defined by the de Boor-Cox recursion formula ^56^:

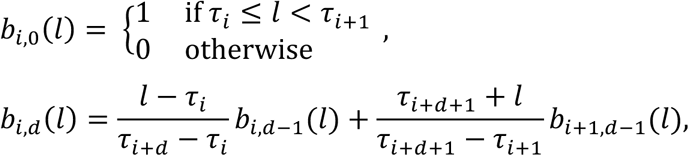

where (𝜏_1_, 𝜏_2_, ⋯, 𝜏*_n_*_+*d*+1_) is a knot vector that divides the parameter space into subintervals, and *n* is the number of basis functions used for the approximation. A continuous surface is approximated by a linear combination of the tensor products of the B-spline basis functions, as follows:

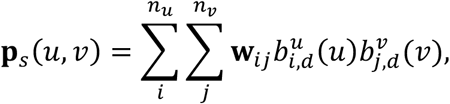

where 𝐰*_ij_*, is a coefficient vector called a control point, and 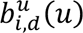 and 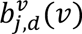 are B-spline basis functions of degree *d* along the independent parameters (*u*, *v*) of the parameter space. In this study, we parameterized the leaf surface by applying a principal component analysis (PCA), and the 2D parameter space was defined using the 1st and the 2nd PC axes. The *𝐰_ij_*, is estimated by minimizing the sum of the squared distances between the B-spline surface and point cloud data. Based on the reconstructed surface, the parameter space is trimmed using a closed B-spline curve to exclude the outer region of the leaf. The closed B-spline curve was fitted to the point cloud data projected into the parameter space using the asymmetric distance with a penalty for concavity and error-adaptive knot insertion (EAKI)^55^. In this study, we used the surface/on_nurbs submodule ^57^ in the Point Cloud Library (PCL; https://pointclouds.org/) to reconstruct leaf surfaces using B-spline surface fitting and B-spline curve trimming. In the B-spline surface fitting, we estimated the 3-degree B-spline surface by incrementally adding knots three times using the following parameter values: the surface smoothness, boundary smoothness, and boundary weight were 0.8, 0.5, and 0.2, respectively. In the B-spline curve trimming, the 3-degree B-spline curve was estimated with a maximum of 100 iterations using the following parameter values for the penalty for concavity (0.2) and for EAKI: iterations without knot insertion and the maximum number of control points were 4 and 100, respectively.

Based on both methods, leaf area was estimated using dataset_v_ through scenario iv. Each leaflet was cropped from the point-cloud data using Cloud Compare v2.12.4 (CloudCompare), software for visualizing and manipulating point-cloud data. We then applied color thresholding (i.e., excluding points with 0.3 < hue < 0.7, saturation < 0.2, and value > 0.8 in the HSV color space) and statistical outlier removal (i.e., excluding outlier points for which the mean distance calculated from their 30 neighbor points is greater than three standard deviations by using the statistical_outlier_removal function in Open3D).

## Results

### Optimizing image magnification for mask image generation

We identified the optimal magnification of input images for multiview images of soybean individuals by evaluating the IoU and processing time of segmentation.

For both three- and four-class segmentations, the IoU increased with decreasing magnification, peaking at approximately 1/4×, followed by a gradual decrease (Fig. 3A, F). The IoU by class showed the same pattern (Fig. 3B-D, G-J); however, several classes peaked at 1/5.4×: stake (three-class segmentation) 0.8670, background (four-class segmentation) 0.9939, and stage (four-class segmentation) 0.9903 (Fig. 3C, H-J). There is a tradeoff: as the magnification of the input images decreases, a single input tile to the U-Net covers larger plant structures; conversely, as the magnification increases, the resolution decreases. In our dataset, a magnification of 1/4× or 1/5.4× reached the optimal IoU.

The processing times for both the three- and four-class segmentations decreased with decreasing magnification. This is because the total number of input tiles decreased (Fig. 2C, 3E, and 3K). Although a marginal increase in processing time was shown to be linked to reductions in the magnification (i.e., the expansion of the field of view), almost all the processing times per image fall between 0.015–0.02 s (Fig. S2).

The U-Net models for the three- and four-class segmentations were trained using the images of all four cultivars in the dataset_t_ (see Fig. S3). These models generated high-resolution mask images for SfM/MVS, in which the background was excluded (Fig. 2D): soybean mask images (soybean pixels were the foreground, whereas others were the background) and soybean/stage mask images (soybean and stage pixels were the foreground, whereas others were the background).

Moreover, we developed a web application for mask generation based on the trained U-Net models. This application helps generate mask images with arbitrary scales and overlapping thresholds via a simple user interface (Fig. 4). The processing time was 126 s on the GPU device for predicting 256 mask images of 5184 × 3456 pixels divided into two columns and two rows of 2900 × 2900 pixel tiles (approximately 1/5.4×) with at least 10% overlap using a four-class segmentation model.

### Quality of reconstructed point cloud data through SfM/MVS with four mask image usages

We reconstructed the dense point cloud data of six individuals of three cultivars at three different growth stages of the dataset_m_ based on SfM/MVS, following four possible scenarios of mask image usage (Fig. 5). Masking during SfM/MVS effectively suppressed the noise derived from the background textures (Fig.5B-D, 5F-H, 5K, and 5L) compared to scenario (i) without mask images (Fig. 5A, 5E, 5I), as noted in previous studies (e.g., ^37,38^).

The quality of the reconstructed point-cloud data varied depending on the mask type and timing of the setting. Although in scenario (ii), setting soybean mask images before SfM was able to suppress the noise derived from the background textures, the reconstructed points along the leaf edges seemed inaccurate (Fig. 5B, 5F) and often failed to reconstruct the leaf surfaces for the small soybean plants (Fig. 5J). These inaccurate reconstructions without background texture-derived noise were a consequence of the low accuracy of camera extrinsic parameter estimation in SfM, not using markers on the stage and corresponding points on the pot, because the ROIs based on the mask images were limited to soybeans. In scenario (ii), key points and tie points were located only in the region corresponding to soybean (Fig. 5N), and their detected numbers were lower than those in the other scenarios (Table 3). There were few qualitative differences between scenarios (iii) and (iv), except for the noise derived from the abaxial side of the leaves and the background textures (Fig. 5C, 5D, 5G, 5H, 5K, and 5L). In both scenarios, leaf surfaces were well-reconstructed because SfM precisely estimated extrinsic camera parameters using the markers on the stage and the corresponding points on the pot.

### Evaluation of the throughput of SfM/MVS on four mask image **usages**

The processing times, including those for mask image generation, SfM, and MVS, were measured for the four scenarios (Fig. 6). Scenarios (ii), (iv), and (iii) achieved the first, second, and third shortest processing times, respectively (.A). The decrease in the area of the ROIs caused by setting mask images contributed to the processing time. The longest mean processing time was 34.0 min for scenario (i), and the shortest one was 15.8 min for scenario (ii).

**Fig 6.**
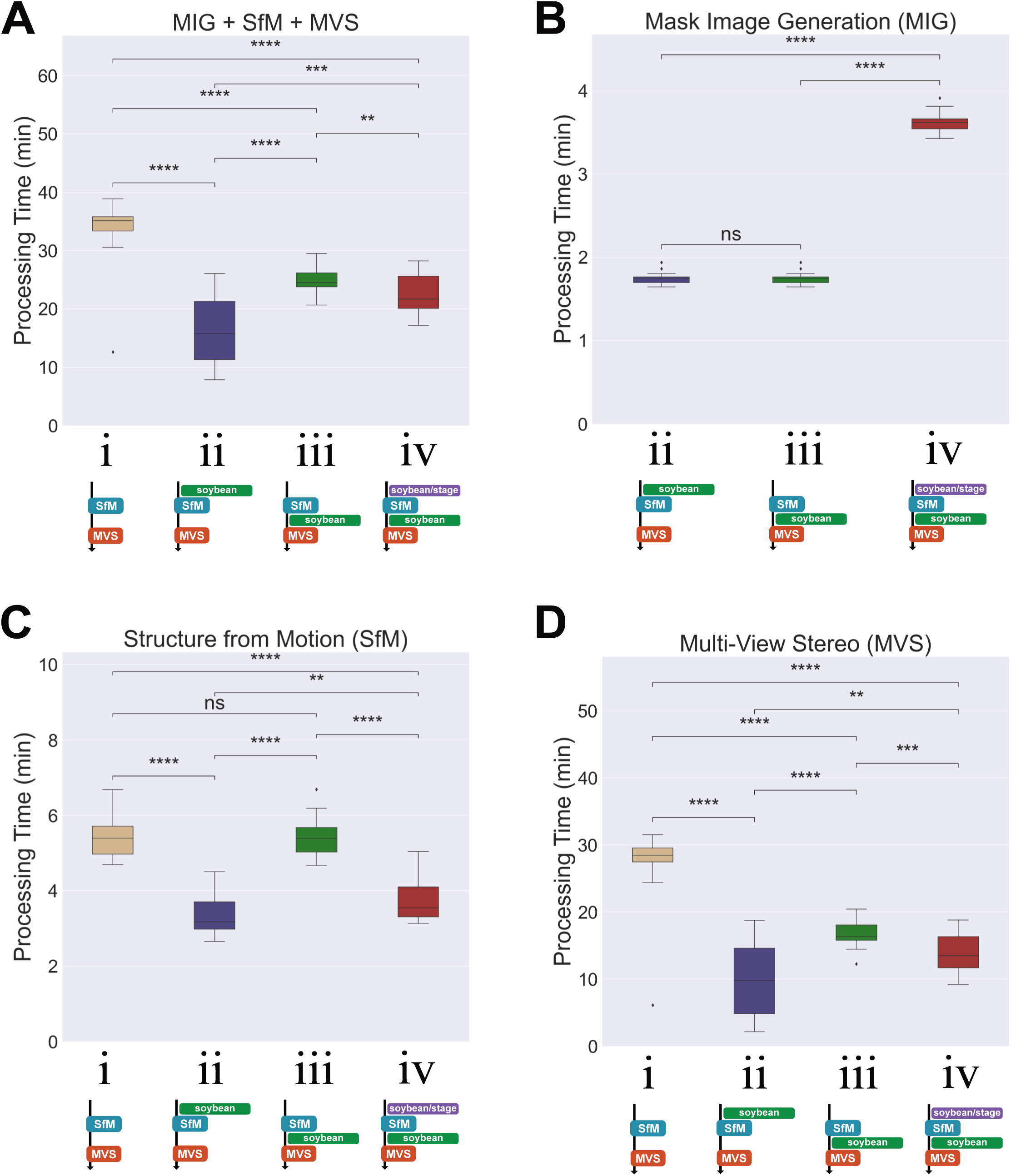
The processing times on the four different mask usage scenarios. The processing times of (A) total of mask image generation (MIG), SfM, and MVS, (B) MIG, (C) SfM, and (D) MVS. Multiple comparison was conducted on the processing times among four scenarios (Mann-Whitney *U* test, **: 0.001<p<=0.01, ***: 0.0001<p<=0.001, ****: p<=0.0001).

The processing time of mask image generation for scenario (iv) was 3.61 min, approximately double that for 1.74 min of scenario (ii) and 1.74 min of scenario (iii) (Fig. 6B), because two types of mask images (soybean and soybean/stage) were generated.

The processing times of SfM and MVS increased in the order of the ROIs, except for scenarios (ii) and (iii) of MVS (Fig. 6C and 6D). There was no significant difference between the processing times of SfM in scenarios (i) and (iii), and the mean processing times were ∼5.4 min. The shortest mean processing time was 3.3 min for scenario (ii). The majority of processing times came from the MVS; the longest mean processing time was 27.6 min in scenario (i) and the shortest one was 9.6 min in scenario (ii). Although the ROIs for MVS were the same between scenarios (ii) and (iii), scenario (ii) showed a shorter processing time for MVS because fewer key points were detected in SfM in scenario (ii) (Table 3).

We adopted scenario (iv) for the following analysis because (1) it generated the highest-quality point cloud data, and (2) its entire processing time was the second shortest.

### Leaf surface reconstruction methods for evaluating phenotypic values

We reconstructed the leaf surfaces of 190 leaflets from dataset_v_, excluding dead leaves, using both Poisson surface reconstruction and B-spline surface fitting, and then estimated their leaf areas (Table S1). The estimated leaf area had the following correlation coefficients with the results of the destructive measurements: Poisson surface reconstruction (𝜌 = 0.9123; Fig. 7A) and B-spline surface fitting (𝜌 = 0.9943; Fig. 7B). The approach based on B-spline surface fitting enabled accurate estimation of leaf area (𝑅^)^ = 0.9871; Fig. 7B). Using the Poisson surface reconstruction, the leaf area was overestimated, particularly for larger leaves. Leaf surface reconstruction based on B-spline surface fitting achieved higher accuracy than Poisson surface reconstruction for the leaf area estimation in almost all cases. Both methods successfully reconstructed leaf surfaces and estimated areas on high-quality point cloud data (Fig. 7C); the Poisson surface reconstruction method enabled a more detailed reconstruction of leaf surfaces, yet showed nearly equivalent performance in leaf area estimation. Although the Poisson surface reconstruction occasionally produced the leaf surface as dual layers for the point cloud data to reconstruct the abaxial side over the adaxial side, the B-spline surface fitting consistently reconstructed a singular surface (Fig. 7D). However, when a single leaflet was reconstructed as multi-leaf-like point cloud data, neither method could accurately reconstruct the surface, resulting in an overestimation of the area (Fig. 7E); we found four leaflets corresponding to this case (Table 4). Three of the four leaflets were leaflets of the same individual plant, which implied blurring associated with the motion of the target plants while capturing the multi-view images, which caused low accuracy of the camera parameter estimation. This is different from the decrease in accuracy in camera parameter estimation owing to the mask images observed for the small individuals in scenario (ii) (Fig. 5J); in fact, the quality of the reconstructed point cloud data is similar across scenarios (ii) and (iv) (Fig. 7F).

**Fig 7.**
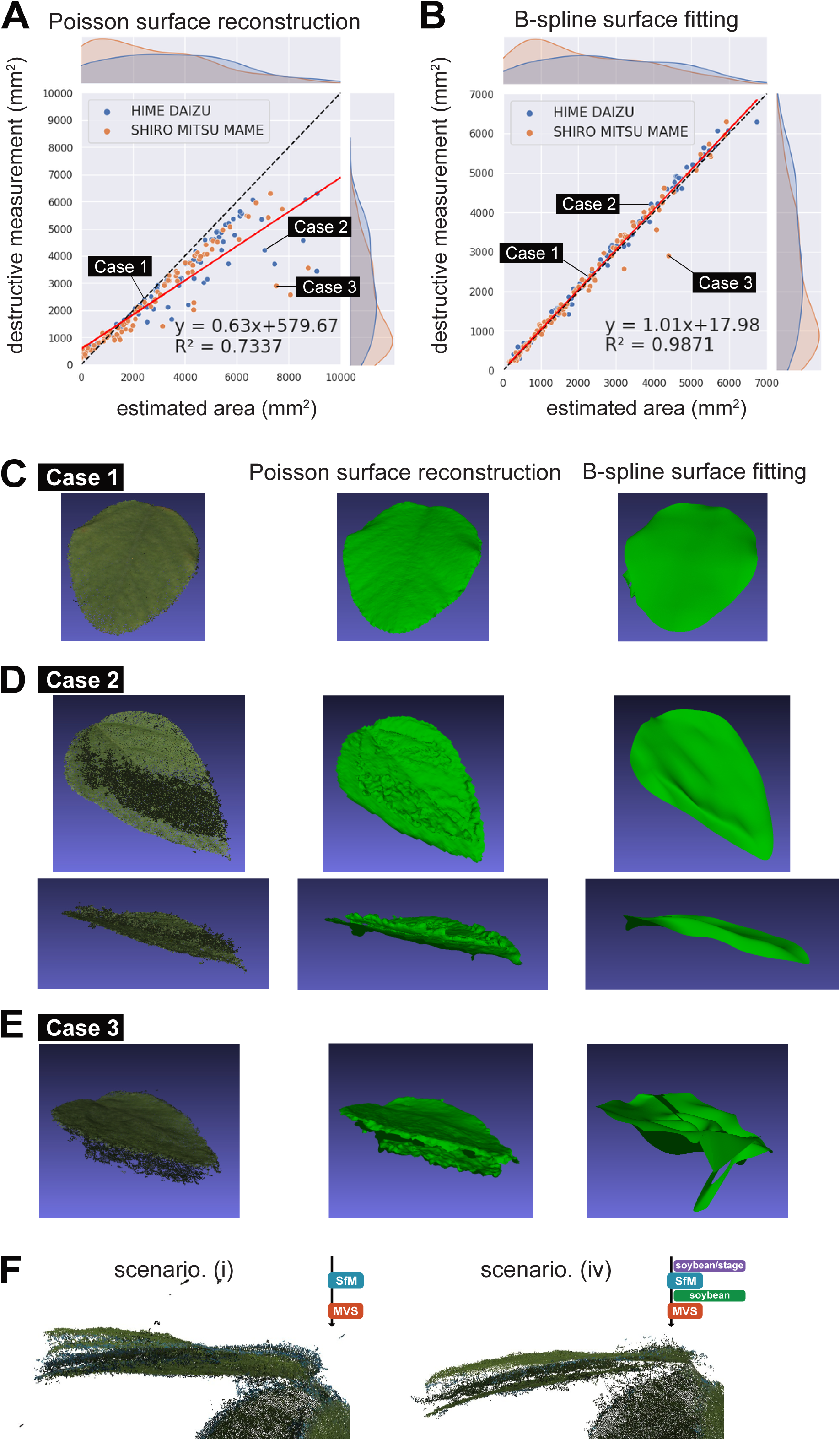
Leaf surface reconstruction and leaf area estimation. Scatter diagrams of leaf area estimated by (A) Poisson surface reconstruction and (B) B-spline surface fitting, against the destructive measurements. (C) Both methods estimated leaf area consistent with the destructive measurements (Hime daizu #7 S01_t). (D) Poisson surface reconstruction overestimated leaf area while B-spline surface fitting estimated accurately (Hime daizu #7 S07_t). (E) Both methods failed to reconstruct leaf surface well, then overestimated leaf area (Shiro mitsu mame #3 S06_r). (F) The reconstructed 3D point cloud data (Shiro mitsu mame #3 S07_t) based on scenario (i) and (iv).

## Discussion

Even though using mask images improves the quality of point cloud data reconstructed via the SfM/MVS pipeline and decreases the processing time, it remains crucial to understand which objects should be masked and in which operations these mask images should be applied. As noted in previous studies ^37,38^, the reconstructed point-cloud data based on SfM/MVS were improved for large individuals by masking the target objects (Fig. 5A-H). However, the point cloud was not comprehensively reconstructed by setting the mask images before SfM for small individuals (Fig. 5J), although they were reconstructed properly without mask images (Fig. 5I). We found that appropriate point clouds were reconstructed either by setting soybean mask images after SfM (scenario iii) or by setting soybean/stage mask images before SfM and then soybean mask images after SfM (scenario iv) (Fig. 5C, D, G, H, K, and L). This implies the importance of setting mask images, which ensures a sufficient number and accuracy of detected key points and tie points (Fig. 5M-P, Table 3). There are several considerations for generating the mask images used in this study, one of which is the magnification rate of the input images. As a result of different scales of input images to the U-Net, a magnification of 5.4× achieved the first or second optimal IoU for all classes (Fig. 3). However, there was a trade-off between low magnification (i.e., long-range context is available with low resolution) and high magnification (i.e., only local context is available with high resolution) (Fig. 2C). Setting a larger input image size could resolve this issue; however, in practice, balancing them for the target objects is necessary because of various constraints (e.g., available GPU memory).

In plant phenotyping, it is advantageous to introduce known morphological constraints to the target shape. In this study, we assumed that a leaf constitutes a single bounded surface in a three-dimensional space during leaf surface reconstruction, which enabled more robust measurements of leaf area (Fig. 7). Thus, the surface was reconstructed using Poisson surface reconstruction; this approach resulted in an overestimation of leaf area (Fig. 7A). It is worth noting that point cloud data derived from actual plants often contain noise that is difficult to eliminate, even with denoising filters. In cases where B-spline surface fitting is ineffective for reconstruction, methods exclusively derived for characterizing leaf morphology could prove useful (e.g., ^39–41,44,45^). The results of leaf area estimation using B-spline surface fitting showed a high correlation with the results of destructive measurements (Fig. 7B), indicating its utility in the phenotyping process. In some cases, the B-spline surface fitting did not reconstruct precisely (Fig. 7E). However, this issue often stems from the difficulty underlying camera parameter estimation in SfM (Fig. 7F), demonstrating that improvements in measurement devices can yield improvements in accuracy. Concerning the qualitatively noted shortcomings in previous studies of the B-spline surface fitting approach in reconstructing leaf surfaces ^44,45^, it is considered that these issues do not significantly impact leaf area estimation as long as the point cloud is reconstructed with high accuracy. Consequently, we propose an optimal workflow for estimating leaf area of soybean plants, including mask image generation using U-Net with 1/5.4× magnification, 3D point cloud reconstruction based on scenario (iv) of SfM/MVS, and leaf surface reconstruction by B-spline surface fitting to estimate leaf area. The optimal values and methods depend on the target objects and constraints on the measurement processes, and the optimal configurations are explored through comparative experiments, as demonstrated in this study. The reconstructed leaf surfaces also allow for quantification of leaf orientation in three-dimensional space; the resultant outputs can be used for functional assessments, including light interception ^58,59^. Utilizing this method in functional structural plant models (FSPM) and other areas is a subject for future research on sustainable food systems using data-driven approaches ^7,8,60^.

Moreover, we offer a tool and data that can be applied toward enhancing the efficiency at which plant phenotyping can be conducted. Specifically, our web application facilitates the generation of mask images through a trained U-Net model, and is designed to allow the replacement of models, enabling its use in various subjects (Fig. 4). We also published the multiview soybean images used in model training as open data, including their mask images and images from destructive measurements. These data will be available to developers of future methodologies as benchmark datasets. We believe that disseminating such tools and datasets will contribute to the field of phenomics, where data acquisition is currently labor-intensive and slow.

## Conclusion

Here, we investigated 3D reconstruction-based phenotyping processes including mask image generation using U-Net models, 3D reconstruction based on the SfM/MVS pipeline, and leaf surface reconstruction for leaf area estimation. Through comparative experiments using soybean data, we proposed an optimal workflow for estimating leaf area and generating mask images. The workflow entailed using U-Net models with 1/5.4× magnification, 3D point cloud reconstruction based on the SfM/MVS pipeline setting soybean/stage mask images before SfM and soybean mask images after SfM (scenario iv), and leaf surface reconstruction by B-spline surface fitting to estimate leaf area. However, it is important to note that in some cases, greater accuracy was achieved with 1/4 magnification in the U-Net model, and that high-quality point clouds can also be obtained by setting only the soybean mask images after SfM (scenario iii). The processes outlined in this study can be adapted as required. Moreover, we offer a web application for mask generation, a command-line tool for leaf surface reconstruction, and validation datasets to facilitate future plant phenotyping research and methodological development.

## Acknowledgements

We thank Horiguchi, R., Inbe, N., Suzuki, M., and Kudo, Y. for their assistance in making multi-view image datasets. We acknowledge the National Agriculture and Food Research Organization (NARO) for providing the seeds used in this study. The Environmental Control Center for Experimental Biology at Kyushu University provides cultivation space.

## Author contributions

KN conceived and designed this study. DS and KN performed the implementation and analyzed the results. DS and KN were major contributors to writing the manuscript. All authors read and approved the final manuscript.

## Data availability

The datasets and source code used during the current study are available in Zenodo (https://doi.org/10.5281/zenodo.10065546, https://doi.org/10.5281/zenodo.10488045, https://doi.org/10.5281/zenodo.10489251, https://doi.org/10.5281/zenodo.10489292, and https://doi.org/10.5281/zenodo.10489326), GitHub (https://github.com/MorphometricsGroup/mask-prediction-app), and DockerHub (https://hub.docker.com/r/noshita/leaf-surf-recon).

## Supplementary Information

**Figure S1. Photogrammetry system.**

**Figure S2. Processing times of mask image generation per single 512×512 tile.**

**Figure S3. Time series data of class-weighted Dice loss and IoU of the U-Net models during training on the entire dataset_t_.**

**Table S1. Leaf area.**

## Figures and Tables

**Table 1. Specimen list in the dataset_m_.**

**Table 2. Cross-validations of the U-Net models for three- and four-class segmentation.** For each fold, different values of class weights were used for calculating class-weighted Dice loss.

**Table 3. The average numbers of key points and tie points.** The number of key points and tie points detected in SfM/MVS pipeline were counted for dataset_m_ across four scenarios.

**Table 4. The four leaflets which were overestimated leaf area using B-spline surface fitting.** The leaf areas of 190 leaflets were overestimated when B-spline surface fitting against the destructive measurements was used.

## References

1. Tilman, D., Balzer, C., Hill, J. & Befort, B. L. Global food demand and the sustainable intensification of agriculture. Proc Natl Acad Sci U S A 108, 20260– 20264 (2011).

2. van Dijk, M., Morley, T., Rau, M. L. & Saghai, Y. A meta-analysis of projected global food demand and population at risk of hunger for the period 2010–2050. Nat Food 2, 494–501 (2021).

3. Wu, W. et al. Global cropping intensity gaps: Increasing food production without cropland expansion. Land use policy 76, 515–525 (2018).

4. Potapov, P. et al. Global maps of cropland extent and change show accelerated cropland expansion in the twenty-first century. Nat Food 3, 19–28 (2022).

5. Zhao, C. et al. Temperature increase reduces global yields of major crops in four independent estimates. Proc Natl Acad Sci U S A 114, 9326–9331 (2017).

6. Wang, X. et al. Emergent constraint on crop yield response to warmer temperature from field experiments. Nat Sustain 3, 908–916 (2020).

7. Herrero, M. et al. Innovation can accelerate the transition towards a sustainable food system. Nat Food 1, 266–272 (2020).

8. Fiorani, F. & Schurr, U. Future Scenarios for Plant Phenotyping. Annu Rev Plant Biol 64, 267–291 (2013).

9. van Etten, J. et al. Data-driven approaches can harness crop diversity to address heterogeneous needs for breeding products. Proc Natl Acad Sci U S A 120, (2023).

10. Coppens, F., Wuyts, N., Inzé, D. & Dhondt, S. Unlocking the potential of plant phenotyping data through integration and data-driven approaches. Curr Opin Syst Biol 4, 58–63 (2017).

11. Goodwin, S., McPherson, J. D. & McCombie, W. R. Coming of age: Ten years of next-generation sequencing technologies. Nat Rev Genet 17, 333–351 (2016).

12. Stark, R., Grzelak, M. & Hadfield, J. RNA sequencing: the teenage years. Nat Rev Genet 20, 631–656 (2019).

13. Sathyamoorthy, S. & Okafor, N. Advances and Challenges in IoT Sensors Data Handling and Processing in Environmental Monitoring Systems. TechRxiv 1–8 (2023) doi:10.36227/techrxiv.24045951.v1.

14. Ullo, S. L. & Sinha, G. R. Advances in Smart Environment Monitoring Systems Using IoT and Sensors. Sensors 20, 3113 (2020).

15. Granier, C. & Tardieu, F. Multi-scale phenotyping of leaf expansion in response to environmental changes: The whole is more than the sum of parts. Plant Cell Environ 32, 1175–1184 (2009).

16. Langstroff, A., Heuermann, M. C., Stahl, A. & Junker, A. Opportunities and limits of controlled-environment plant phenotyping for climate response traits. Theoretical and Applied Genetics vol. 135 Preprint at 10.1007/s00122-021-03892-1 (2022).

17. Araus, J. L. & Cairns, J. E. Field high-throughput phenotyping: The new crop breeding frontier. Trends Plant Sci 19, 52–61 (2014).

18. Yang, W. et al. Crop Phenomics and High-Throughput Phenotyping: Past Decades, Current Challenges, and Future Perspectives. Mol Plant 13, 187–214 (2020).

19. Campbell, Z. C., Acosta-Gamboa, L. M., Nepal, N. & Lorence, A. Engineering plants for tomorrow: how high-throughput phenotyping is contributing to the development of better crops. Phytochemistry Reviews 17, 1329–1343 (2018).

20. Dammer, K.-H., Wollny, J. & Giebel, A. Estimation of the Leaf Area Index in cereal crops for variable rate fungicide spraying. European Journal of Agronomy 28, 351– 360 (2008).

21. Addai, I. K. & Alimiyawo, M. Graphical determination of leaf area index and its relationship with growth and yield parameters of Sorghum (Sorghum bicolor L. Moench) as affected by fertilizer application. Journal of Agronomy 14, 272–278 (2015).

22. Paulus, S., Behmann, J., Mahlein, A.-K., Plümer, L. & Kuhlmann, H. Low-Cost 3D Systems: Suitable Tools for Plant Phenotyping. Sensors 14, 3001–3018 (2014).

23. Lin, Y. LiDAR: An important tool for next-generation phenotyping technology of high potential for plant phenomics? Comput Electron Agric 119, 61–73 (2015).

24. Chéné, Y. et al. On the use of depth camera for 3D phenotyping of entire plants. Comput Electron Agric 82, 122–127 (2012).

25. Milella, A., Marani, R., Petitti, A. & Reina, G. In-field high throughput grapevine phenotyping with a consumer-grade depth camera. Comput Electron Agric 156, 293–306 (2019).

26. An, N. et al. Plant high-throughput phenotyping using photogrammetry and imaging techniques to measure leaf length and rosette area. Comput Electron Agric 127, 376– 394 (2016).

27. Santos, T. T. & Rodrigues, G. C. Flexible three-dimensional modeling of plants using low-resolution cameras and visual odometry. Mach Vis Appl 27, 695–707 (2016).

28. Schonberger, J. L. & Frahm, J.-M. Structure-from-Motion Revisited. in 2016 IEEE Conference on Computer Vision and Pattern Recognition (CVPR) vols 2016-Decem 4104–4113 (IEEE, 2016).

29. Snavely, N., Seitz, S. M. & Szeliski, R. Modeling the World from Internet Photo Collections. Int J Comput Vis 80, 189–210 (2008).

30. Seitz, S. M., Curless, B., Diebel, J., Scharstein, D. & Szeliski, R. A Comparison and Evaluation of Multi-View Stereo Reconstruction Algorithms. http://vision.middlebury.edu/mview. (2006).

31. Schönberger, J. L., Zheng, E., Frahm, J. M. & Pollefeys, M. Pixelwise view selection for unstructured multi-view stereo. in Lecture Notes in Computer Science (including subseries Lecture Notes in Artificial Intelligence and Lecture Notes in Bioinformatics) (eds. Leibe, B., Matas, J., Sebe, N. & Welling, M.) vol. 9907 LNCS 501–518 (Springer International Publishing, Cham, 2016).

32. Cernea, D. OpenMVS: Multi-View Stereo Reconstruction Library. Preprint at (2020).

33. Lu, X. et al. Reconstruction method and optimum range of camera-shooting angle for 3D plant modeling using a multi-camera photography system. Plant Methods 16, 118 (2020).

34. Zermas, D., Morellas, V., Mulla, D. & Papanikolopoulos, N. Estimating the Leaf Area Index of crops through the evaluation of 3D models. in IEEE International Conference on Intelligent Robots and Systems vols 2017-Septe 6155–6162 (IEEE, 2017).

35. Koutsoudis, A., Ioannakis, G., Vidmar, B., Arnaoutoglou, F. & Chamzas, C. Using noise function-based patterns to enhance photogrammetric 3D reconstruction performance of featureless surfaces. J Cult Herit 16, 664–670 (2015).

36. Zhao, J., Monno, Y. & Okutomi, M. Polarimetric Multi-View Inverse Rendering. IEEE Trans Pattern Anal Mach Intell 45, 8798–8812 (2023).

37. Nooruddin, M. & Rahman, M. Improved 3D reconstruction for images having moving object using semantic image segmentation and binary masking. in 4th International Conference on Electrical Engineering and Information and Communication Technology, iCEEiCT 2018 32–37 (IEEE, 2018). doi:10.1109/CEEICT.2018.8628064.

38. Tanabata, T., Isobe, S., Hayashi, A. & Kochi, N. Development of a semi-automatic 3D modeling system for phenotyping morphological traits in plants. in Proceedings: IECON 2018 - 44th Annual Conference of the IEEE Industrial Electronics Society 5439–5444 (IEEE, 2018). doi:10.1109/IECON.2018.8592904.

39. Dornbusch, T., Wernecke, P. & Diepenbrock, W. A method to extract morphological traits of plant organs from 3D point clouds as a database for an architectural plant model. Ecol Modell 200, 119–129 (2007).

40. Pound, M. P., French, A. P., Murchie, E. H. & Pridmore, T. P. Automated recovery of three-dimensional models of plant shoots from multiple color images. Plant Physiol 166, 1688–1698 (2014).

41. Kempthorne, D. M. et al. Surface reconstruction of wheat leaf morphology from three-dimensional scanned data. Functional Plant Biology 42, 444–451 (2015).

42. Kazhdan, M., Bolitho, M. & Hoppe, H. Poisson Surface Reconstruction. Eurographics Symposium on Geometry Processing (2006).

43. Yang, M. Der, Chao, C. F., Huang, K. S., Lu, L. Y. & Chen, Y. P. Image-based 3D scene reconstruction and exploration in augmented reality. Autom Constr 33, 48–60 (2013).

44. Zhu, F. et al. 3D Reconstruction of Plant Leaves for High-Throughput Phenotyping. in Proceedings - 2018 IEEE International Conference on Big Data, Big Data 2018 4285–4293 (IEEE, 2018). doi:10.1109/BigData.2018.8622428.

45. Ando, R., Ozasa, Y. & Guo, W. Robust Surface Reconstruction of Plant Leaves from 3D Point Clouds. Plant Phenomics 2021, (2021).

46. Ronneberger, O., Fischer, P. & Brox, T. U-net: Convolutional networks for biomedical image segmentation. in Lecture Notes in Computer Science (including subseries Lecture Notes in Artificial Intelligence and Lecture Notes in Bioinformatics) vol. 9351 234–241 (2015).

47. He, K., Zhang, X., Ren, S. & Sun, J. Deep Residual Learning for Image Recognition. in 2016 IEEE Conference on Computer Vision and Pattern Recognition (CVPR) vols 2016-Decem 770–778 (IEEE, 2016).

48. Buslaev, A. et al. Albumentations: Fast and flexible image augmentations. Information (Switzerland) 11, (2020).

49. Lowe, D. G. Distinctive image features from scale-invariant keypoints. Int J Comput Vis 60, 91–110 (2004).

50. Rothermel, M., Wenzel, K., Fritsch, D. & Haala, N. SURE - Photogrammetric Surface Reconstruction from Imagery. in Proceedings LC3D Workshop (Berlin, 2012).

51. Furukawa, Y., Curless, B., Seitz, S. M. & Szeliski, R. Towards internet-scale multi-view stereo. in Proceedings of the IEEE Computer Society Conference on Computer Vision and Pattern Recognition 1434–1441 (IEEE, 2010). doi:10.1109/CVPR.2010.5539802.

52. Hirschmuller, H. Stereo Processing by Semiglobal Matching and Mutual Information. IEEE Trans Pattern Anal Mach Intell 30, 328–341 (2008).

53. Remondino, F., Spera, M. G., Nocerino, E., Menna, F. & Nex, F. State of the art in high density image matching. The Photogrammetric Record 29, 144–166 (2014).

54. Zhou, Q.-Y., Park, J. & Koltun, V. Open3D: A Modern Library for 3D Data Processing. (2018).

55. Mörwald, T., Balzer, J. & Vincze, M. Modeling connected regions in arbitrary planar point clouds by robust B-spline approximation. Rob Auton Syst 76, 141–151 (2016).

56. de Boor, C. On calculating with B-splines. J Approx Theory 6, 50–62 (1972).

57. Mörwald, T. Object modelling for cognitive robotics. (Technische Universität Wien, 2013).

58. Pearcy, R. W., Muraoka, H. & Valladares, F. Crown architecture in sun and shade environments: assessing function and trade-offs with a three-dimensional simulation model. New Phytologist 166, 791–800 (2005).

59. Niinemets, Ü. Photosynthesis and resource distribution through plant canopies. Plant Cell Environ 30, 1052–1071 (2007).

60. Evers, J. B., Letort, V., Renton, M. & Kang, M. Computational botany: Advancing plant science through functional-structural plant modelling. Ann Bot 121, 767–772 (2018).

