## Supplementary Figures S1-S3. for "A comparative study of plant phenotyping workflows based on three-dimensional reconstruction from multi-view images"

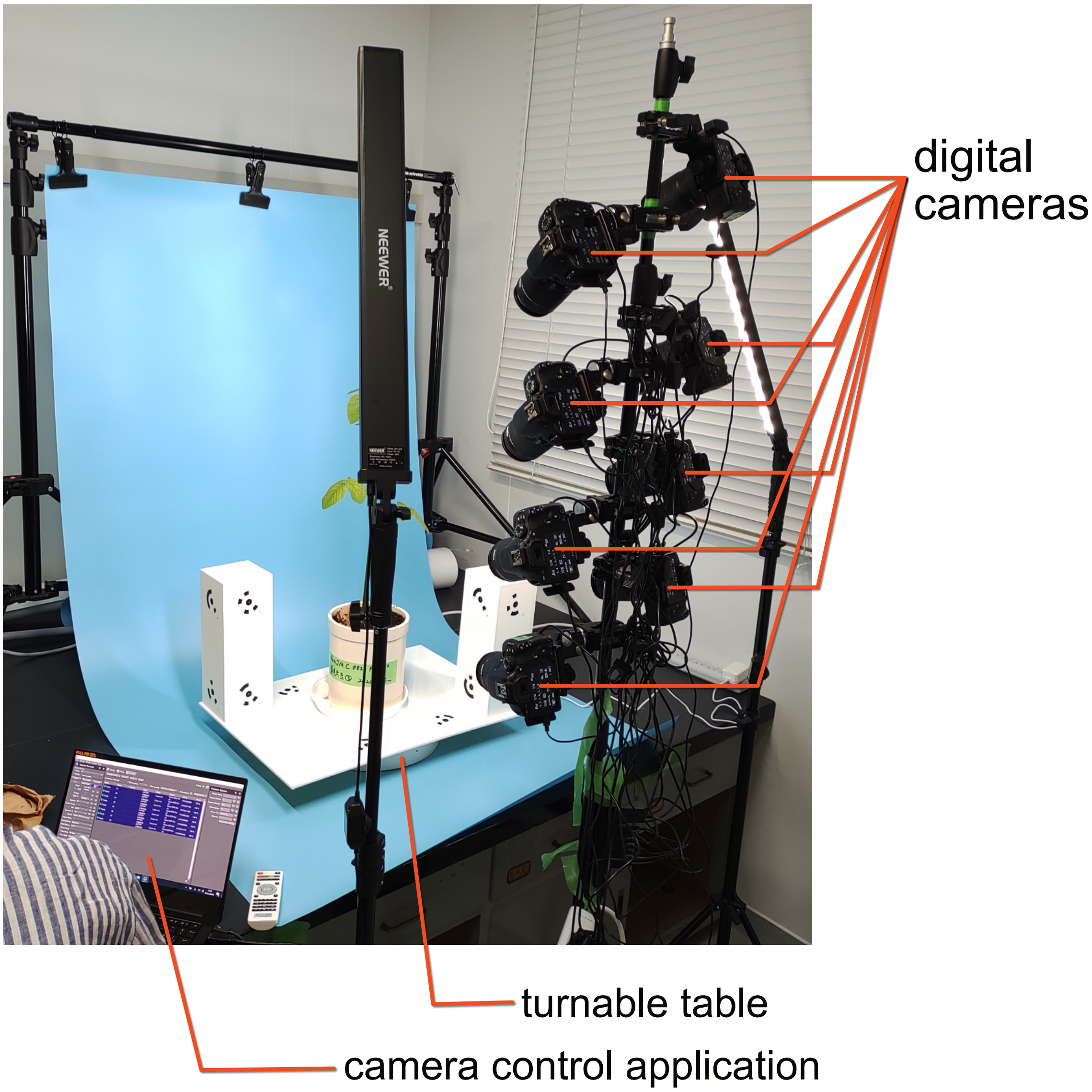


**Figure S1. Photogrammetry system.** This figure was created based on (Noshita, 2023) (Licensed under CC BY 4.0).

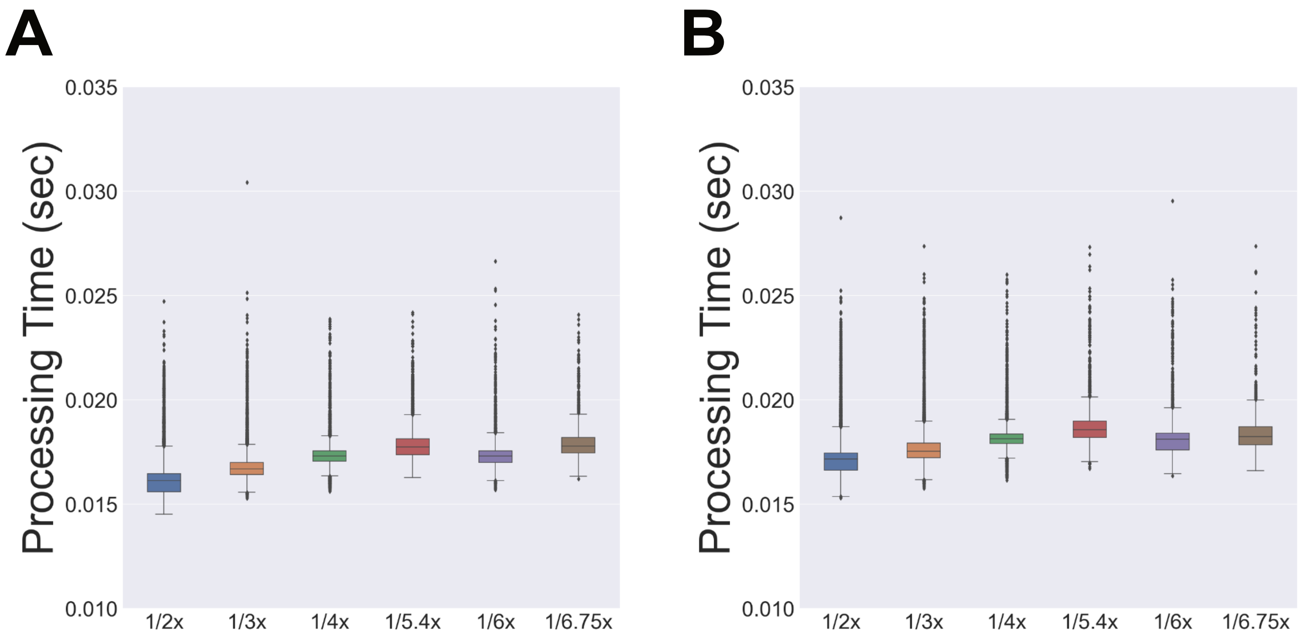


**Figure S2. Processing times of mask image generation per single 512×512 tile.** Box plots of processing times among six magnifications of (a) three-class and (b) four-class segmentation.

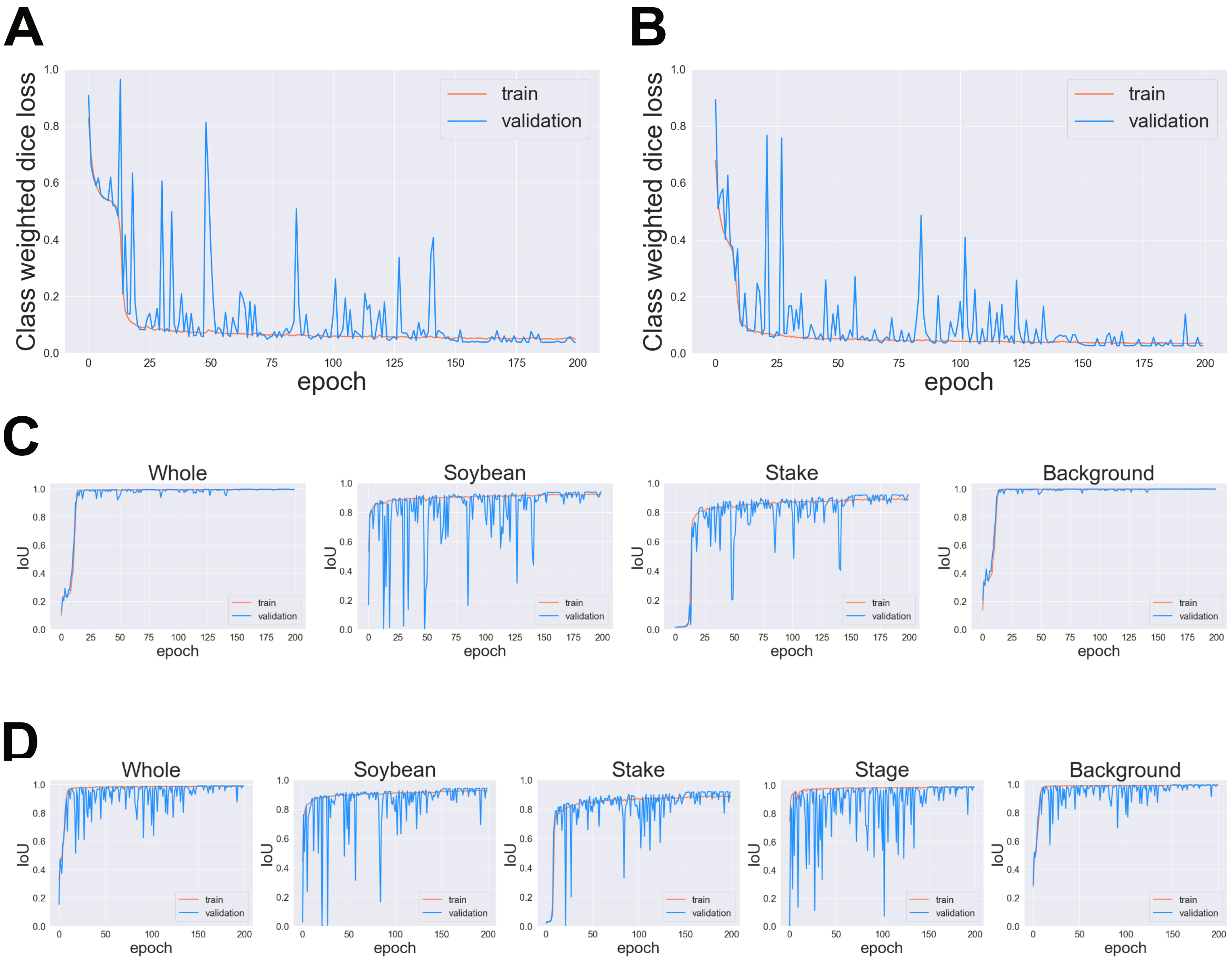


**Figure S3. Time series data of class-weighted Dice loss and IoU of the U-Net models during training on the entire dataset$_t$.** Line plots of class-weighted Dice losses of (a) three-class and (b) four-class segmentation models. Line plots of IoU scores of (c) three-class and (d) four-class segmentation.
